# Quantitative analysis of the ubiquitin-proteasome system under proteolytic and folding stressors

**DOI:** 10.1101/780676

**Authors:** Jeremy J. Work, Onn Brandman

## Abstract

Aging, disease, and environmental stressors are associated with failures in the ubiquitin-proteasome system (UPS), yet a quantitative understanding of how stressors affect the proteome and how the UPS responds is lacking. Here we assessed UPS performance and adaptability in yeast under stressors using quantitative measurements of misfolded substrate stability and stress-dependent UPS regulation by the transcription factor Rpn4. We found that impairing degradation rates (proteolytic stress) and generating misfolded proteins (folding stress) elicited distinct effects on the proteome and on UPS adaptation. Folding stressors stabilized proteins via aggregation rather than overburdening the proteasome, as occurred under proteolytic stress. Still, the UPS productively adapted to both stressors using separate mechanisms: proteolytic stressors caused Rpn4 stabilization while folding stressors increased *RPN4* transcription. In some cases, adaptation completely prevented loss of UPS substrate degradation. Our work reveals the distinct effects of proteotoxic stressors and the versatility of cells in adapting the UPS.

## Introduction

The ubiquitin-proteasome system (UPS) is the primary route for the disposal of defective proteins in eukaryotic cells (Hershko et al. 1983, 1984; Lecker, Goldberg, and Mitch 2006). Aging, genetic mutations, and environmental changes all challenge the UPS and can lead to accumulation of defective proteins (“proteotoxic stress”), which is a hallmark of many neurodegenerative diseases, including Alzheimer’s disease, Parkinson’s disease, Huntington’s disease, and amyotrophic lateral sclerosis (Labbadia and Morimoto 2015; Sweeney et al. 2017; Klaips, Jayaraj, and Hartl 2018). Characterizing the performance and adaptability of the UPS in clearing defective proteins under proteotoxic stressors is thus likely to aid in understanding numerous diseases.

In the UPS, ubiquitin ligases modify selected proteins with polyubiquitin chains that target them for degradation by the 26S proteasome, a 2.5 MDa protein complex composed of 33 unique subunits (Voges, Zwickl, and Baumeister 1999). To simultaneously and stoichiometrically drive the expression of dozens of proteasomal components along with other UPS-related genes, eukaryotes have evolved master transcriptional regulators that target all proteasome genes, as well as ubiquitin, ubiquitin ligases, and extrinsic proteasome factors (Mannhaupt et al. 1999; Lundgren et al. 2003; Meiners et al. 2003; Kraft et al. 2006; Xu et al. 2008; Sato et al. 2009; Radhakrishnan et al. 2010). In budding yeast, this master regulation occurs via the transcription factor Rpn4 (Xie and Varshavsky 2001).

To adapt the expression of UPS components based on cellular needs, cells regulate Rpn4 levels via multiple stress-sensitive mechanisms. These include proteasomal degradation of Rpn4 via two encoded degradation signals (degrons) that target it to the proteasome: one ubiquitin-independent signal at the N-terminus, and one signal recognized by the E3 ubiquitin ligase Ubr2 (Ha, Ju, and Xie 2012; L. Wang et al. 2004). Due to these degrons, Rpn4 has a short half-life of 2 minutes and will therefore quickly accumulate if the proteasome is impaired (Xie and Varshavsky 2001). Additionally, *RPN4* is transcriptionally regulated by several stress-sensitive transcription factors, including Yap1, a responder to oxidative stress; Pdr1/3, the drivers of the pleiotropic drug resistance response; and Hsf1, the driver of the heat shock response (HSR)(Hahn, Neef, and Thiele 2006; Ma and Liu 2010; Temple, Perrone, and Dawes 2005; Moye-Rowley 2003). We term the collective, stress-responsive transcriptional regulation of the UPS through Rpn4 the <u>p</u>roteasome <u>s</u>tress response (“PSR”), analogous to terminology used to describe other transcriptional stress responses like the HSR.

While the PSR has been demonstrated to respond to proteotoxic stressors (Xie and Varshavsky 2001; X. Wang et al. 2010; Schmidt et al. 2019), quantification of its effectiveness at combating such stressors and the relative contributions of its distinct activation mechanisms have not been investigated in diverse proteotoxic conditions. Two ways stressors may increase levels of misfolded proteins are to 1) cause proteins to misfold or obstruct their folding (“folding stress”), or 2) impair degradation rates of misfolded proteins (“proteolytic stress”) (Figure 1A). A naive expectation is that folding and proteolytic stressors have overlapping effects on the proteome and UPS. For example, misfolded proteins generated by a folding stressor may become targeted to the proteasome, increasing competition between proteasome substrates and thereby lowering degradation rates for each substrate (i.e. a folding stressor leading indirectly to proteolytic stress). Conversely, UPS substrates that are stabilized by a proteolytic stressor may potentiate the misfolding of other proteins (i.e. a proteolytic stressor indirectly leading to folding stress), as has been observed when expression of one misfolded protein causes others to misfold (Satyal et al. 2000; Gidalevitz et al. 2006, 2009). However, these hypotheses have not yet been quantitatively evaluated. Furthermore, it is unknown if activation of the PSR fully neutralizes proteotoxic challenges (“perfect adaptation”), if cells accumulate defective proteins in spite of PSR activation (“partial adaptation”), if cells overreact to these challenges (“overadaptation”), and whether cellular responses are distinct for proteolytic and folding stressors.

**Figure 1:**
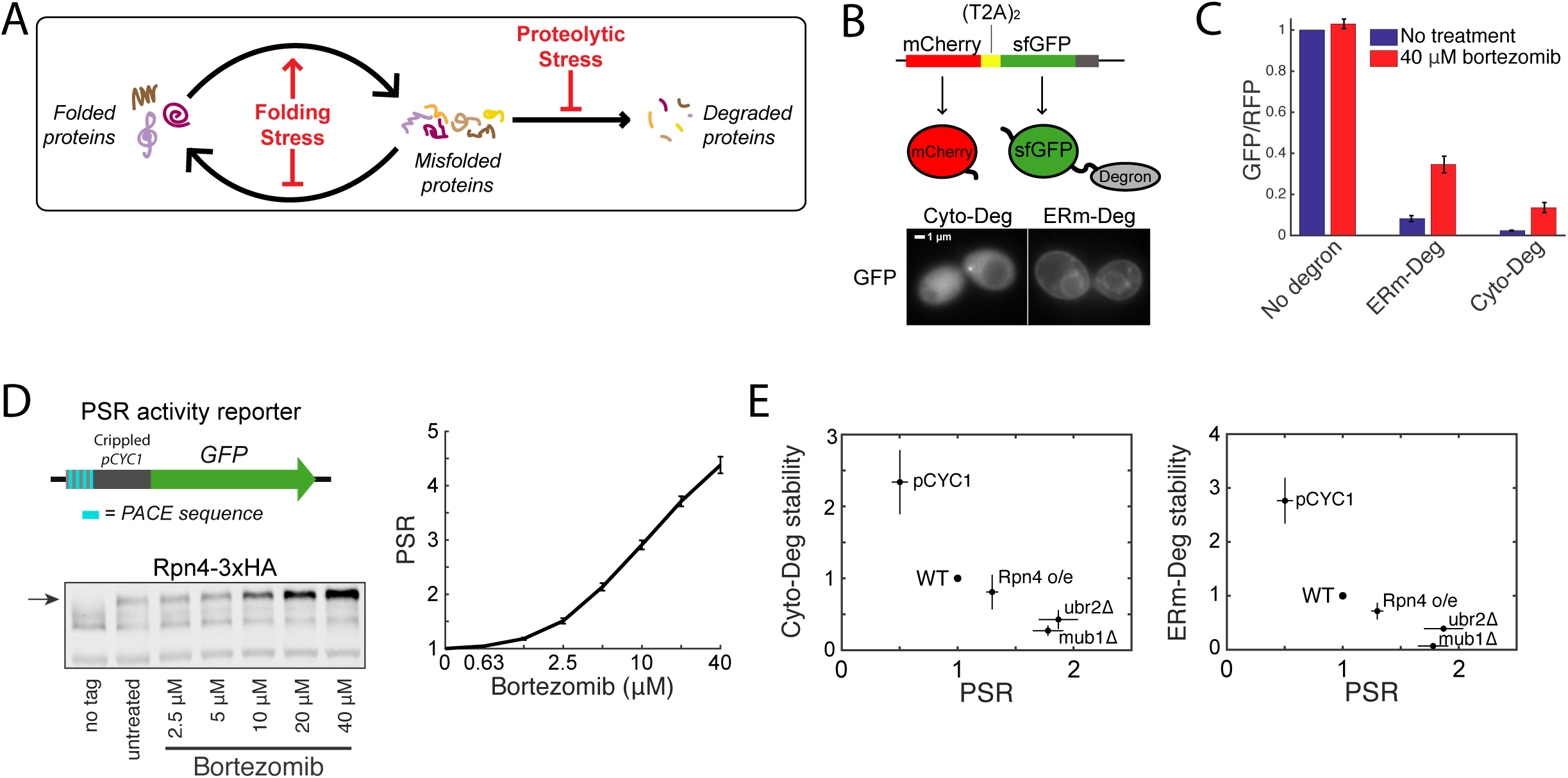
Clearance of defective proteins scales with the PSR. **A:** Diagram of protein folding and degradation, and the effect of folding and proteasomal stress. **B:** (Top) Schematic of T2A system for controlled expression of degron reporters. (Bottom) GFP localization of Cyto-Deg and ERm-Deg in normal conditions. **C:** Mean RFP-normalized GFP fluorescence of Cyto-Deg, ERm-Deg, and a no degron control with either 0 μM (blue bars) or 40 μM (red bars) bortezomib. **D:** (Top left) Schematic of PSR activity reporter. (Bottom left) Immunoblot of HA-tagged endogenous Rpn4 under a serial titration of bortezomib. Arrow indicates the position of Rpn4-3xHA. (Right) Mean forward scatter normalized GFP of PSR activity reporter under a serial titration of bortezomib. Units are fold change from no treatment. **E:** Plots of fold degron stability (Left panel: Cyto-Deg; Right panel: ERm-Deg) versus fold PSR upon modifying Rpn4 levels through deletion of *UBR2* or *MUB1*, replacement of the endogenous *RPN4* promoter with *pCYC1*, or expression of a second copy of *RPN4* from a plasmid. Error bars denote standard error for n≥3 biological replicates.

We systematically characterized the performance and adaptability of the UPS under diverse stress conditions in yeast using quantitative measurements of UPS performance (measured by the stability of misfolded reporter substrates) and PSR-mediated adaptation (transcriptional activation of Rpn4 target genes as measured by a synthetic reporter). We unexpectedly found that proteolytic and protein folding stressors stabilized misfolded proteins and activated the PSR through separate, non-overlapping mechanisms. Proteasomal inhibition (a proteolytic stressor) blocked degradation of misfolded proteins and stabilized Rpn4 without increasing *RPN4* transcription. By contrast, the addition of the amino acid analogs canavanine and AZC (folding stressors) caused aggregation of misfolded proteins rather than their targeting to the proteasome, yet still activated the PSR by driving transcriptional activation of *RPN4* without increasing Rpn4 stability. The PSR productively responded to both proteolytic and protein folding stressors despite their different underlying mechanisms for increasing misfolded protein levels. In both cases, this included perfect or near-perfect adaptation. Our work reveals the adaptability of the UPS and provides a framework to quantitatively understand how cells regulate the UPS in response to proteotoxicity and disease-causing states.

## Results

### Clearance of defective proteins scales with the PSR

To investigate the UPS under stress conditions, we built quantitative reporters that measure the performance of the UPS and the activity of the cell’s primary transcriptional effector of the UPS, the PSR. Because a major role of the UPS is to identify and degrade defective proteins, we defined “UPS performance” as the ability to degrade reporter proteins containing constitutively misfolded domains. We designed two UPS substrates, Cyto-Deg and ERm-Deg, consisting of a green fluorescent protein (sfGFP) fused to a degron sequence featuring a cluster of hydrophobic residues (Maurer et al. 2016)(Figure 1B). Cyto-Deg localizes to the cytosol and is dependent on Hsp70 for degradation (Maurer et al. 2016). ERm-Deg localizes to the endoplasmic reticulum (ER) membrane—likely because its C-terminal degron is also an ER targeting signal—and is ubiquitylated by the ER-membrane-localized ubiquitin ligase Doa10 (Maurer et al. 2016). Doa10 is part of the ERAD-C pathway, which targets ER transmembrane proteins with a misfolded domain in the cytosol (Ruggiano, Foresti, and Carvalho 2014). Thus, expression of both Cyto-Deg and ERm-Deg leads to misfolded protein domains in the cytosol, but they are efficiently degraded by the proteasome via separate pathways. Evaluating the stability of both degrons therefore informs on the state of the proteasome, the shared feature in their degradation. To control for protein synthesis rate, we expressed a red fluorescent protein (mCherry) upstream of the sfGFP-degron fusion, separated by two tandem T2A peptide-skipping sequences (Donnelly et al. 2001; Szymczak and Vignali 2005). mCherry and the sfGFP-degron are synthesized stoichiometrically, but mCherry is detached during translation and escapes UPS targeting. The sfGFP/mCherry ratio is therefore proportional to the stability of the degron-fused protein, where a high ratio indicates high stability and a low ratio indicates low stability. Cyto-Deg and ERm-Deg were both capable of reporting on UPS performance, as evidenced by an increase in the sfGFP/mCherry ratio upon treatment with a 40 µM dose of the proteasome inhibiting drug bortezomib (Figure 1C).

To measure the PSR, we built a synthetic promoter specifically sensitive to changes in the PSR. The PSR is driven by the binding of Rpn4 to a DNA motif called the proteasome-associated control element (PACE), which is found in the promoters of all proteasomal subunits and many proteasome-associated factors (Mannhaupt et al. 1999; Shirozu, Yashiroda, and Murata 2015). The PSR reporter features four tandem copies of the PACE sequence along with a minimal promoter to drive expression of sfGFP (Figure 1D). We validated the reporter’s sensitivity by blocking Rpn4 degradation with bortezomib and observing a monotonic increase in GFP expression in response (Figure 1D). This increase corresponded to increased levels of hemagglutinin-tagged endogenous Rpn4 (Figure 1D).

We first used our reporters to evaluate how modulating the PSR affects UPS performance in unstressed conditions by genetically altering the constitutive levels of Rpn4. To reduce Rpn4 levels, we replaced the endogenous *RPN4* promoter with a weaker promoter (*pCYC1*). To increase Rpn4 levels, we deleted factors necessary for Rpn4 degradation (Ubr2 or Mub1)(Ju et al. 2008; L. Wang et al. 2004), or expressed a second copy of *RPN4* from a single-copy plasmid. PSR activity was inversely correlated with Cyto-Deg and ERm-Deg stability, demonstrating that PSR activity is tightly coupled to UPS performance (Figure 1E).

### Proteotoxic stressors elicit multiple adaptive regimes

To understand the adaptive potential of the UPS, we investigated the role of the PSR in clearing defective proteins during proteotoxic stress. We achieved this by measuring the PSR and degron stability in cells after 5 hour treatment with three proteotoxic compounds: bortezomib, canavanine (an arginine analog), and azetidine-2-carboxylic acid (“AZC”; a proline analog). Bortezomib directly inhibits the proteasome to cause proteolytic stress. Canavanine and AZC directly disrupt protein folding when incorporated into newly synthesized proteins, causing folding stress. By measuring cellular responses after 5 hours, we aimed to capture the system after adaptive mechanisms had taken effect. At the highest concentrations tested, all three stressors increased the stability of both Cyto-Deg and ERm-Deg and induced the PSR, consistent with their proteotoxicity (Figure 2A). We were concerned that high doses of canavanine and AZC would directly disrupt PSR reporter inducibility, so we limited the concentrations of canavanine and AZC to a range in which the PSR reporter remained comparably inducible by 5 µM bortezomib and could therefore reliably report on PSR activity (Figure 2B).

**Figure 2:**
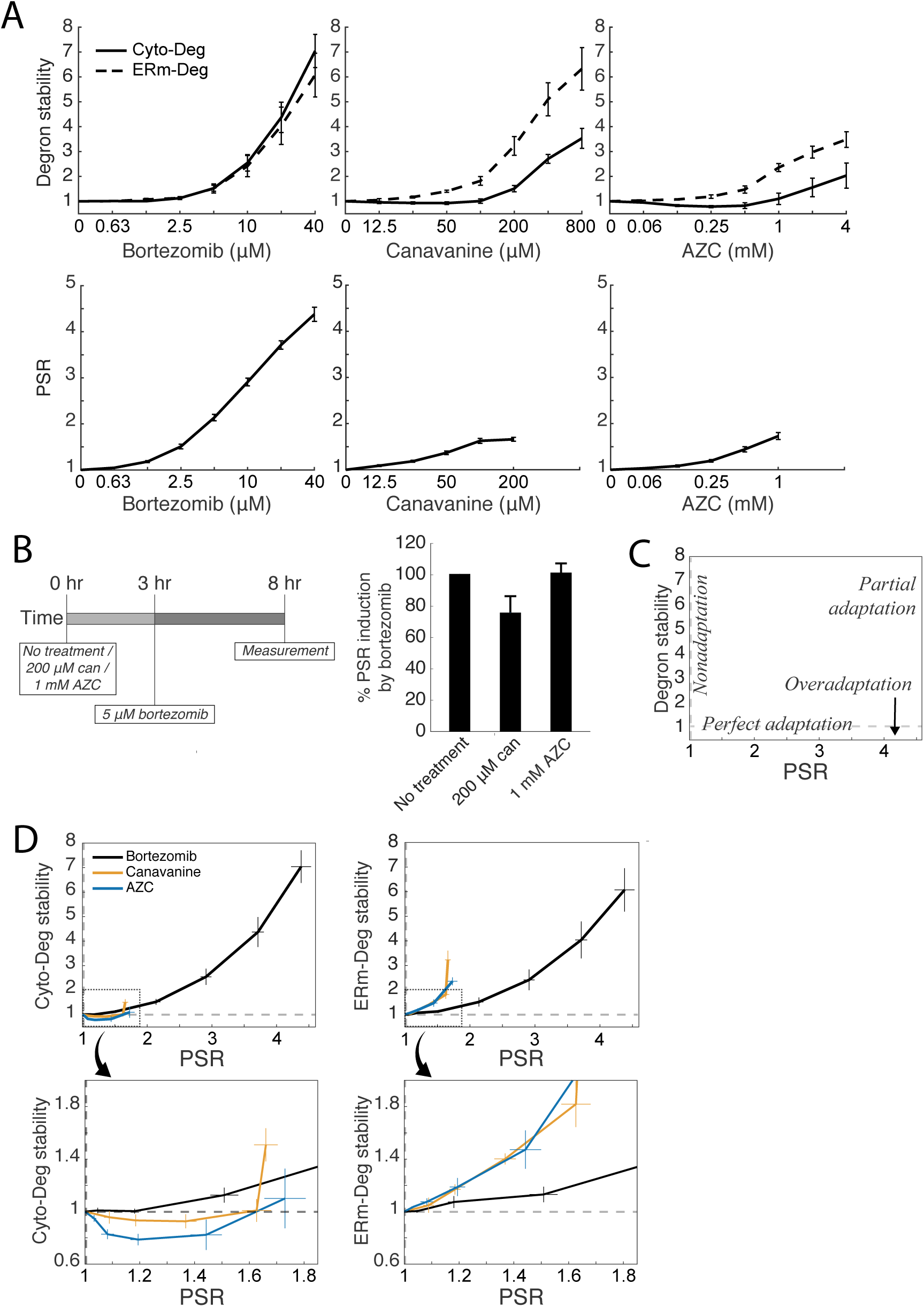
Proteotoxic stressors elicit multiple adaptive regimes. **A:** Measurements of fold degron stability (top row, solid: Cyto-Deg, dotted: ERm-Deg) and fold PSR activity (bottom row) in titrations of bortezomib, canavanine, or AZC. **B:** Measurement of PSR activity (right) after treatment with 5 μM bortezomib and either 200 μM canavanine, 1 mM AZC, or no additional treatment, at the noted time points (left). **C:** Schematic of adaptive regimes as a function of degron stability and PSR activity. **D:** Plots of fold degron stability (left: Cyto-Deg, right: ERm-Deg) versus fold PSR activity for the titrations in (A) to reveal the adaptive regime for each stressor. The boxed regions in the upper plots are enlarged in the lower plots. Error bars denote standard error for n≥3 biological replicates.

To determine how the UPS adapts to each stressor, we compared the relationship between UPS performance and PSR activation (Figure 2C). *Perfect adaptation*, a regime where cells respond to a stressor without any loss of UPS performance, would be observed as activation of the PSR without any change in UPS performance. *Non-adaptation* would manifest as a lack of PSR activation with concurrent loss of UPS performance. *Partial adaptation* would present as an intermediate between these two regimes, where the PSR activates but UPS performance still declines. Finally, *overadaptation* would be evidenced by PSR activation coupled to an increase in UPS performance. Strikingly, cells exhibited near-perfect adaptation in response to low doses of bortezomib (≤2.5 µM), as the PSR was activated but stability remained the same or nominally increased for both degrons (Figure 2A,2D). At higher doses (>2.5 µM), the response to bortezomib resulted in partial adaptation, showing decreasing UPS performance as bortezomib dose increased despite PSR activation. Responses to canavanine and AZC caused a divergent response: UPS adaptation was perfect or overadaptive up to 200 µM and 1 mM respectively for Cyto-Deg but partial for ERm-Deg. These observations suggest UPS adaptation is highly effective but becomes less so under severe stressors, as noted by an increase in stability of both degrons at high doses of bortezomib and ERm-Deg at several doses of canavanine and AZC. Furthermore, the divergence of adaptive response between proteolytic stressors (partial adaptation regardless of degron monitored) and folding stressors (degron-specific adaptation regimes) suggests that these stressors cause distinct effects on cells.

### Activating the PSR improves UPS performance under stress

To understand the limitations of the PSR, we next investigated why adaptation was imperfect for ERm-Deg under AZC and canavanine treatment and for both degrons at high doses of bortezomib. One model to explain these results is that the PSR is insufficiently activated in these conditions, resulting in a PSR that cannot fully compensate for the increased proteotoxic burden. Alternatively, the stressors we applied may exceed the capacity of the PSR to increase UPS performance. To distinguish between these models, we tested whether enhancing the PSR by expressing a second copy of *RPN4* improves degradation in stress conditions. Indeed, it was recently shown that *RPN4* overexpression improves UPS performance in clearing mislocalized ER proteins or defective ribosome proteins in the cytosol (Schmidt et al. 2019; Tye et al. 2019). As expected, expressing a second copy of *RPN4* in the absence of stress increased the PSR and destabilized Cyto-Deg and ERm-Deg relative to an empty vector control (Figure 1E and 3). Under stress conditions, the addition of a second *RPN4* copy lowered degron stability for nearly all concentrations of bortezomib, canavanine, or AZC tested (Figure 3). We conclude that activating the PSR is sufficient to improve UPS performance under all stressors and that Rpn4 is insufficiently activated to clear specific substrates in partially adaptive regimes.

**Figure 3:**
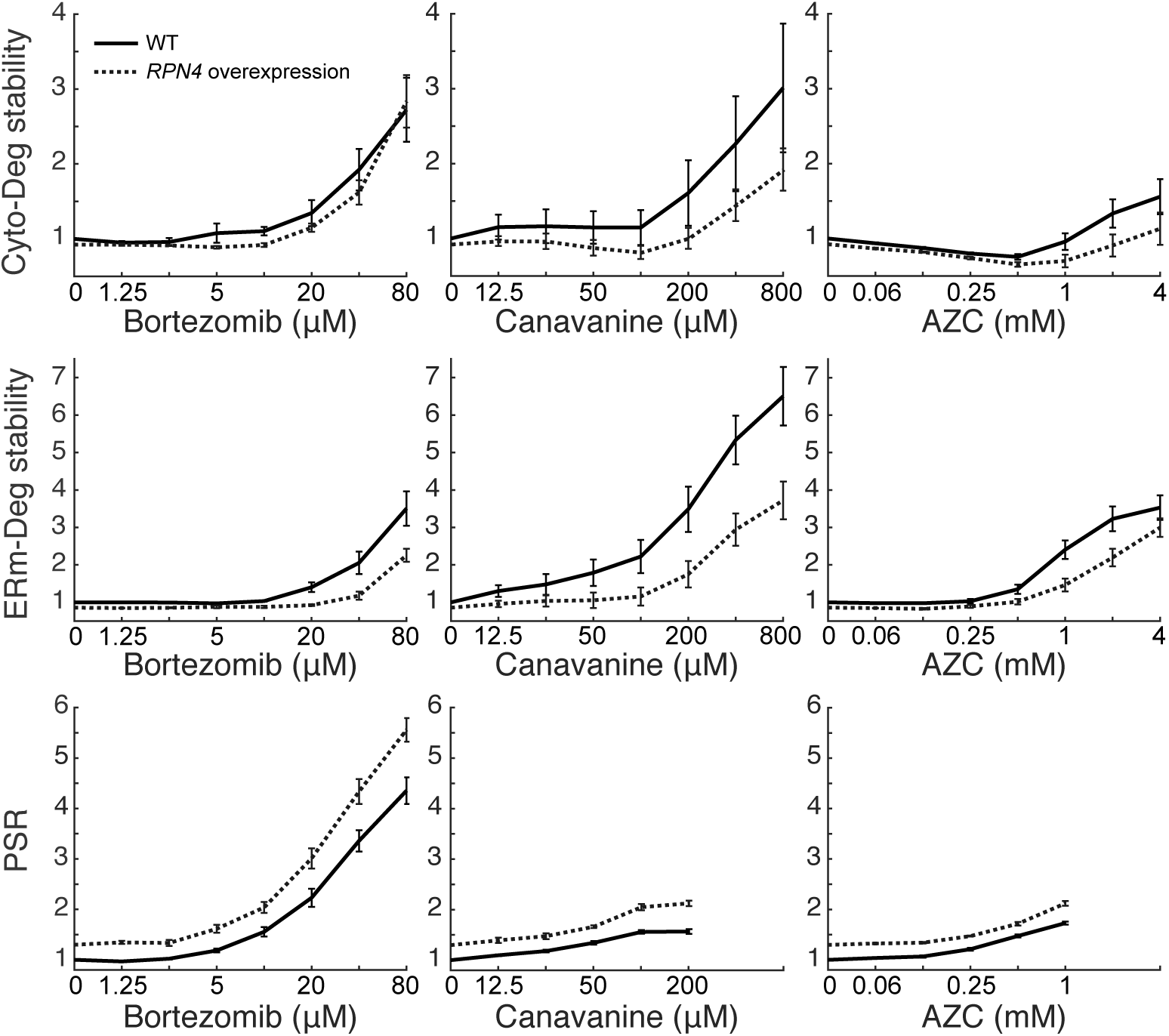
Boosting the PSR improves UPS performance under stress. Measurements of fold degron stability (top: Cyto-Deg, middle: ERm-Deg) and fold PSR activity (bottom) in titrations of bortezomib, canavanine, or AZC for cells expressing an empty vector (black) or a second copy of *RPN4* (dotted). Error bars denote standard error for n≥3 biological replicates.

### Folding stressors activate the PSR predominantly via transcription of *RPN4*

Because amino acid analogs caused a degron-specific adaptive regime while bortezomib did not (Figure 2D), we reasoned that these stressors may be sensed differently by cells. Proteasome inhibition via bortezomib is likely to activate the PSR by impaired degradation of Rpn4, although it may also cause transcriptional activation of *RPN4*. In contrast, it is unclear whether the folding stressors canavanine and AZC activate the PSR through creation of new proteasome substrates that compete with Rpn4 for degradation and stabilize it (i.e. an indirect proteolytic stress), transcriptional activation of *RPN4*, or both. Indeed, canavanine and AZC robustly activate Hsf1, a transcriptional activator of *RPN4* (Hahn, Neef, and Thiele 2006; Yamamoto et al. 2008; Alford and Brandman 2018). We therefore investigated the role of transcriptional regulation of *RPN4* in activating the PSR during stressor treatment. Consistent with previous work, a reporter of Hsf1 activity (Brandman et al., 2012) robustly activated in the presence of canavanine and AZC (up to 6- and 7-fold activation at their highest concentrations, respectively), with comparatively weak (up to 2-fold) maximal activation by bortezomib (Figure 4A). Because Hsf1 targets the *RPN4* promoter (Hahn, Neef, and Thiele 2006), we predicted that the promoter of RPN4 (*pRPN4)* should be upregulated in canavanine and AZC stress. We measured the activity of *pRPN4* using a *pRPN4:GFP* plasmid reporter and found that bortezomib did not activate *pRPN4*, while canavanine and AZC modestly increased the promoter’s activity (1.6- and 1.3-fold)(Figure 4B). This is consistent with a model that folding stressors but not proteolytic stressors upregulate the PSR via transcriptional regulation of *RPN4*.

**Figure 4:**
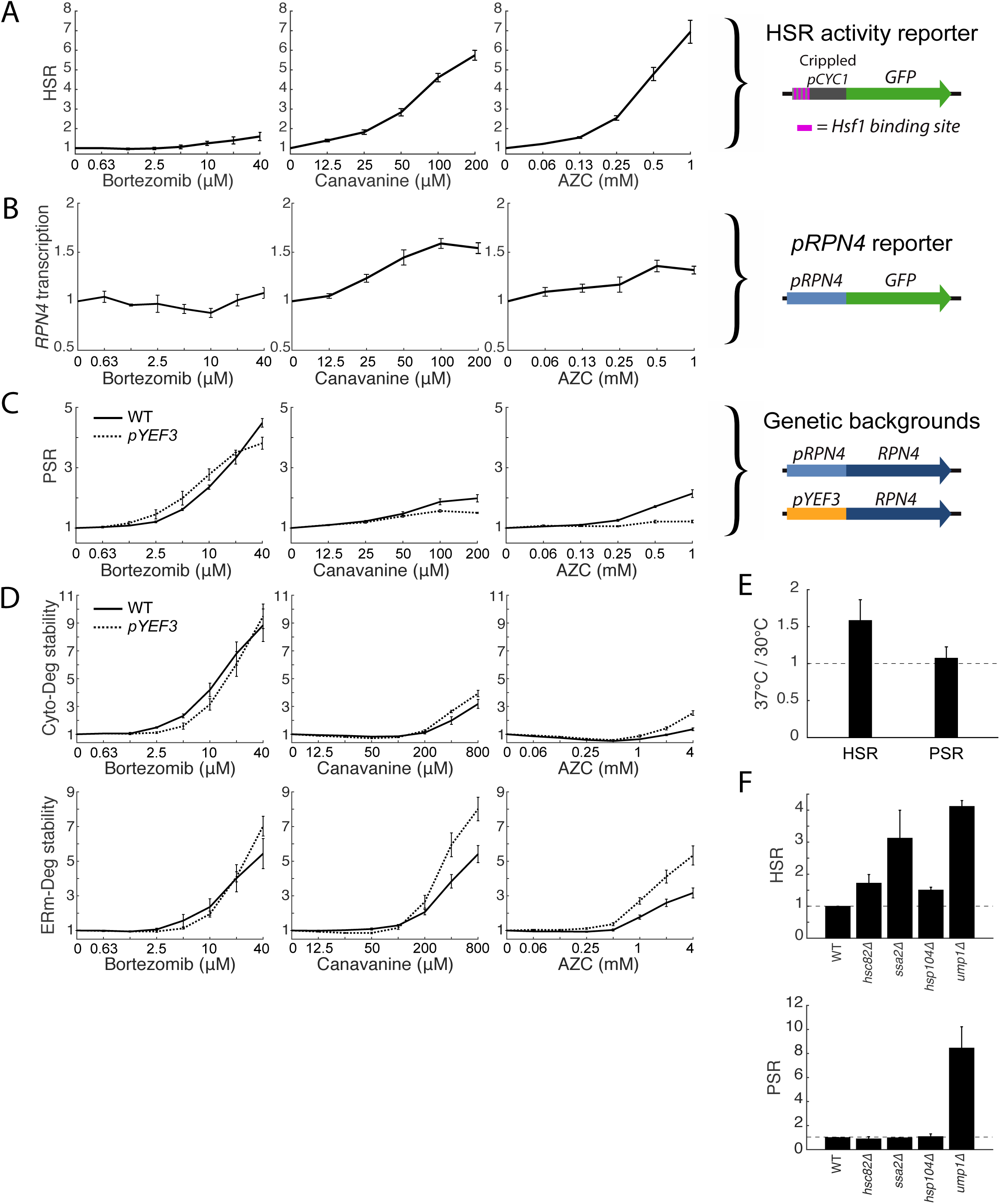
Folding stressors activate the PSR via transcription of RPN4 and do not increase UPS substrate load. **A:** Schematic of a HSR reporter (right) and its fold GFP induction in titrations of bortezomib, canavanine, and AZC (left). **B:** Schematic of a pRPN4 reporter (right) and its fold GFP induction in titrations of bortezomib, canavanine, and AZC (left). **C:** Schematic of the RPN4 locus in wildtype or a pYEF3:RPN4 background (right), and measurements of fold PSR activity in stressor titrations for both strains (left). **D:** Measurements of Cyto-Deg (top) and ERm-Deg (bottom) fold stability for titrations of bortezomib, canavanine, or AZC in either a wildtype (solid) or pYEF3:RPN4 (dotted) background. Titrations of both strains are normalized to the no-treatment values of each reporter. **E:** Mean fold activity of the HSR or PSR at 37°C relative to 30°C in wildtype cells. **F:** Mean fold activity of the HSR (left) or PSR (right) relative to wildtype upon deletion of HSC82, SSA2, HSP104, or UMP1. Error bars denote standard error for n≥3 biological replicates.

### Folding stressors do not increase UPS substrate load

Given the similarity in magnitude of PSR activation and *pRPN4* induction in canavanine and AZC (Figure 2A and 4B), we hypothesized that PSR activation by these two stressors is fully accounted for by transcriptional targeting of *pRPN4*, with little or no contribution through stabilization of Rpn4. By contrast, bortezomib does not activate *pRPN4* (Figure 4B), and presumably induces the PSR through stabilization of Rpn4 alone. To determine the contribution of Rpn4 stabilization to the PSR under canavanine and AZC treatments, we disabled transcriptional regulation by engineering strains in which the genomic copy of *RPN4* is under the control of the *YEF3* promoter (*pYEF3*), which is not targeted by Hsf1 but retains the same approximate basal expression (1.23 fold basal, s.e.=.027). In the absence of stress-sensitive transcriptional regulation of *RPN4*, PSR induction was retained in bortezomib treatment, but partially lost in canavanine and fully lost in AZC treatment (Figure 4C). We therefore predicted that loss of promoter-mediated PSR activation will impair adaptation to folding stressors and minimally affect proteolytic stressors. Indeed, Cyto-Deg and ERm-Deg stability was sensitized to canavanine and AZC in the *pYEF3:RPN4* background, evidenced by greater degron stabilization relative to wildtype (Figure 4D). Furthermore, degron stabilization was greater under AZC treatment than canavanine treatment, which correspondingly had a greater loss of PSR activation in the *pYEF3:RPN4* background. We conclude that protein folding stress caused by canavanine and AZC does not lead to stabilization of Rpn4, and instead UPS adaptation under these stressors occurs predominantly via increased *RPN4* transcription.

Transcriptional activation of the PSR by canavanine and AZC (Figure 4C) could be specific to these treatments or it could be the general response to folding stressors. To test the generality of this phenomenon, we increased the likelihood of protein misfolding by raising the steady state temperature of wildtype yeast from 30°C to 37°C. This was sufficient to activate the HSR, but caused no change in PSR activation (Figure 4E). Consistent with a model in which folding stress does not increase the burden of substrates on the proteasome, deletion of protein chaperone genes (*HSC82, SSA2, HSP104*) increased the HSR but not the PSR (Figure 4F). These results suggest that proteins that misfold due to folding stressors do not create proteolytic stress.

### Folding stressors cause aggregation and result in failure to target aggregation-prone substrates to the proteasome

Given that the misfolded proteins generated by folding stressors did not appear to increase proteasome burden, we explored the possibility that the misfolded proteins are instead sequestered into aggregates in which they are protected from proteasomal degradation. Indeed, it has been previously reported that AZC can cause aggregation of endogenous proteins (Weids and Grant 2014), and that aggregation can be a mechanism for avoiding degradation (Wallace et al. 2015). Because canavanine and AZC robustly activate the HSR, a response that is driven by a drop in protein chaperone availability (Zheng et al. 2016; Alford and Brandman 2018), we reasoned that chaperones become limiting under folding stressors and this may drive protein aggregation. To test this, we boosted Hsf1 activity to increase chaperone levels and determined if this would increase the stability of our degron reporters during canavanine and AZC treatment. We expressed an extra copy of Hsf1 with an N-terminal truncation that renders it constitutively active (Hsf1_Δ1-147_) (Sorger 1990) in a strain expressing *RPN4* from the *CYC1* promoter (to eliminate the confounding effect that Hsf1 activates *pRPN4*). Hsf1_Δ1-147_ reduced Cyto-Deg and ERm-Deg levels in response to canavanine and AZC but not bortezomib (Figure 5A). These results suggest that the decrease in UPS performance due to canavanine and AZC is resolvable by increasing chaperone levels, supporting that they cause protein folding stress, while bortezomib does not. This chaperone dependence for degradation is consistent with the hypothesis that folding stressors result in sequestration of UPS substrates into aggregates.

**Figure 5:**
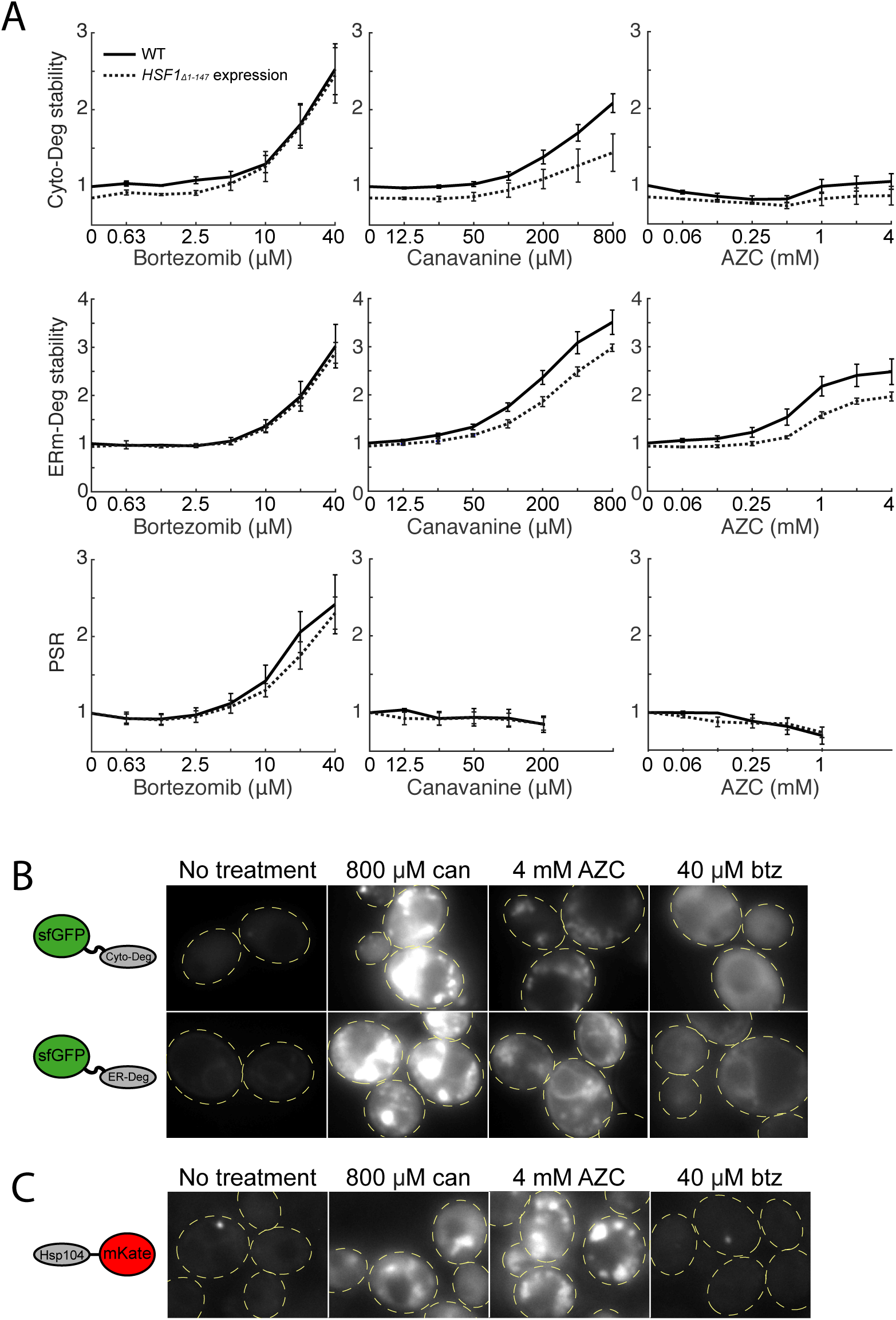
Folding stressors cause aggregation and result in failure to target aggregation-prone substrates to the proteasome. **A:** Measurements of fold degron stability (top: Cyto-Deg, middle: ERm-Deg) and fold PSR activity (bottom) in titrations of bortezomib, canavanine, or AZC for cells expressing an empty vector (solid) or a copy of HSF1Δ 1-147 (dotted). **B-C:** Fluorescent localization of ERm-Deg (GFP), Cyto-Deg (GFP), or Hsp104-mKate2 (RFP) in cells treated with 800 μM canavanine, 4 mM AZC, 40 μM bortezomib, or no treatment for 5 hours. Cells are outlined in yellow dashed lines.

To directly assess the presence of aggregates in canavanine and AZC treated cells, we performed fluorescence microscopy on cells expressing Cyto-Deg or ERm-Deg. Canavanine and AZC caused GFP in both Cyto-Deg and ERm-Deg to form inclusions (Figure 5B). Consistent with previous reports (Weids and Grant 2014), 4 mM AZC also increased expression of Hsp104 and relocalized it into foci (Figure 5C) in cells without a degron reporter. A similar response was observed with 800 µM canavanine. This change in Hsp104 localization suggests that endogenous proteins (not just our synthetic reporters) are sequestered into aggregates in the presence of canavanine and AZC. By contrast, even a high dose (40 µM) of bortezomib that strongly increased degron levels did not alter the localization of the degrons or induce Hsp104 expression. These observations suggest that canavanine and AZC cause protein folding stress that drives misfolded proteins into aggregates rather than targeting them to the proteasome. Under this model, folding stress fails to stabilize Rpn4 because it does not produce proteasome substrates that compete with Rpn4 for degradation.

If aggregation interferes with the degradation of misfolded proteins by the UPS, we predicted that highly soluble proteasome substrates that escape aggregation during folding stress would continue to be degraded normally. To test this prediction, we built a third degron reporter, ATA-Deg (a “CAT tail” degron), which was demonstrated in previous work to be poly-ubiquitylated and targeted for proteasomal degradation, but whose six-peptide degron sequence (“ATAATA”) is soluble (Sitron and Brandman, 2019). Accordingly, ATA-Deg was diffuse throughout the cytosol even under high doses of canavanine, AZC, or bortezomib (Figure 6A). Bortezomib treatment increased the stability of ATA-Deg, indicating sensitivity to proteolytic stress. Conversely, ATA-Deg levels were constant or decreased under all concentrations of canavanine and AZC treatment (Figure 6B), suggesting that folding stress does not impair degradation of ATA-Deg. Consistent with this and in contrast to Cyto-Deg and ERm-Deg (Figure 5A), ATA-Deg stability was not affected by expression of Hsf1_Δ1-147_ under folding stressors. We conclude that folding stressors sequester aggregation-prone UPS substrates without disrupting degradation of soluble substrates (Figure 6D).

**Figure 6:**
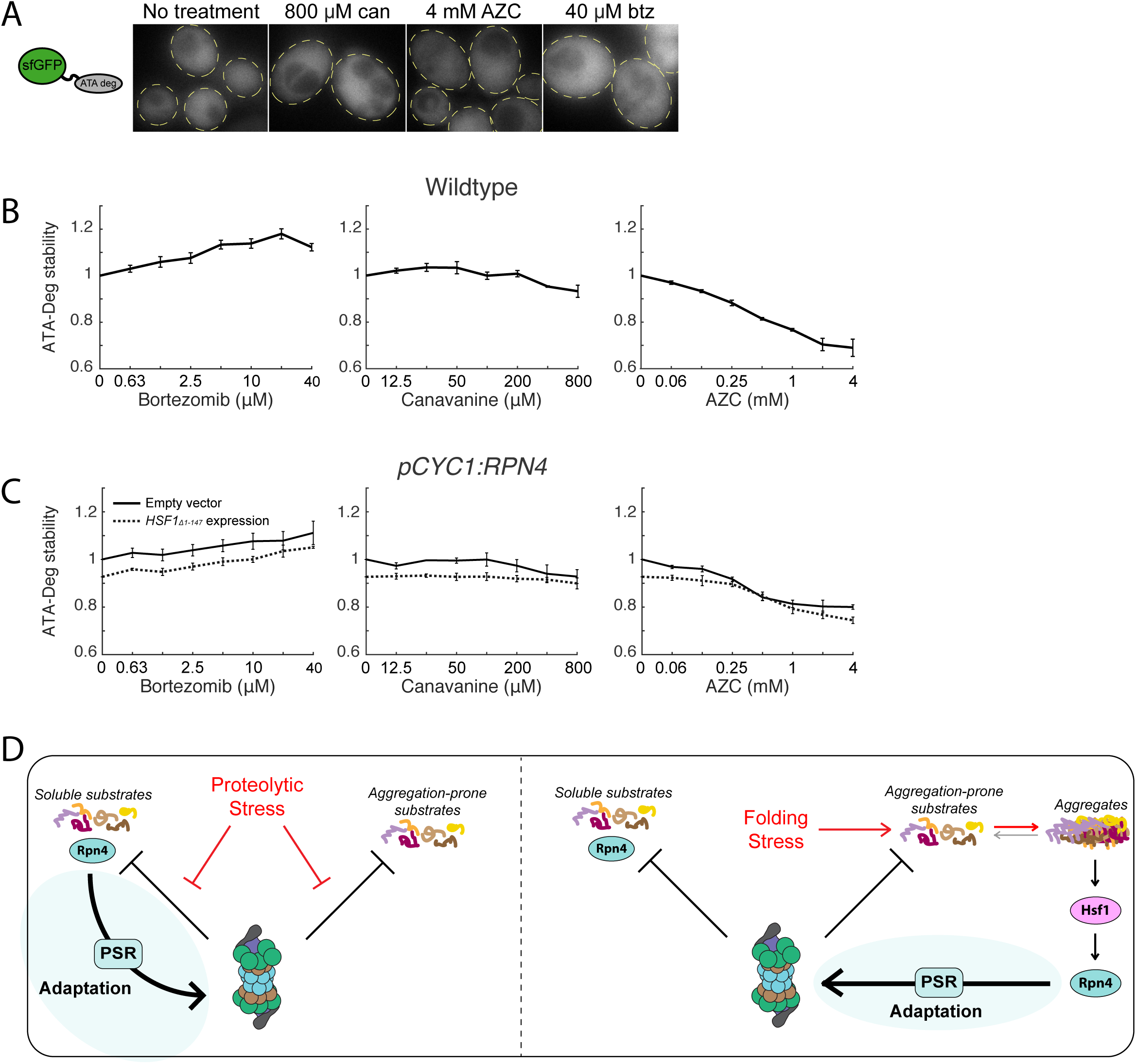
Folding stressors do not impair degradation of soluble UPS substrates. **A:** Fluorescent localization of ATA-Deg (GFP) in cells treated with 800 μM canavanine, 4 mM AZC, 40 μM bortezomib, or no treatment for 5 hours. Cells are outlined in yellow dashed lines. **B:** Measurements of fold ATA-Deg stability in titrations of bortezomib, canavanine, or AZC. **C:** Fold ATA-Deg stability in drug titrations in cells expressing an empty vector (solid) or a copy of HSF1Δ 1-147 (dotted). Error bars throughout denote standard error for n≥3 biological replicates. D: Model for UPS adaptation by the PSR to proteolytic and folding stressors. (Left) Proteolytic stressors increases the stability of all proteasome substrates. This includes Rpn4, whose accumulation leads to PSR activation. (Right) Folding stressors causes some proteasome substrates to sequester into aggregates. Aggregation triggers the HSR, activates transcription of RPN4, and causes PSR activation.

## Discussion

Here we assessed the performance and adaptability of the UPS in yeast under stress conditions using quantitative measurements of UPS performance and the adaptive transcriptional response of the UPS (the “proteasome stress response,” or PSR). We found that proteolytic and protein folding stressors stabilized misfolded proteins through separate, non-overlapping mechanisms, with the former blocking degradation of misfolded proteins and the latter resulting in their aggregation rather than their targeting to the proteasome. Despite a difference in the underlying proteostasis defect, the UPS productively responded to both proteolytic and folding stressors, and in both cases, this included perfect or near-perfect adaptation (no loss in degradation performance) for some substrates (Figure 2D).

The perfect and near-perfect adaptation we observed for the UPS implies the existence of an underlying network that can mechanistically achieve this (Ferrell 2016). In the case of folding stress, *RPN4* is activated transcriptionally (Figure 4B and 4C), likely by Hsf1, to achieve perfect adaptation for proteins that aggregate in the cytosol. This intervention may cause increased degradation rates of soluble proteasome substrates, an intriguing consequence of a system that tunes the UPS to “problem” proteins that are poor UPS substrates (aggregated proteins). This substrate-specific adaptation likely occurs to some degree in all stress responses that use concerted transcriptional regulation to address substrates with distinct adaptive needs.

Under proteolytic stress, where Rpn4 is stabilized and its activation rescues the degradation of other proteasomal substrates (Figure 2D), perfect adaptation requires that the increase in Rpn4 activity compensate for the loss in degradation of proteasome substrates. This may occur via a combination of the multiple degrons present on Rpn4 and could also involve post-translational regulation of Rpn4, which is ubiquitylated and phosphorylated (Ju et al. 2007; L. Wang et al. 2004). Additionally, proteasome inhibition may cause the buildup of substrates that selectively outcompete Rpn4 but not other substrates, making Rpn4 levels hypersensitive to proteolytic stress. Understanding the range of substrates and conditions for which perfect or near-perfect adaptation occurs and its underlying mechanisms is an important topic for future study.

Our data suggest that protein folding stressors do not burden the proteasome with increased overall substrate load (Figure 4C, 4F, and 4G). Instead, misfolding causes the sequestration of aggregation-prone substrates into inclusions and a resultant loss of their UPS targeting. This cellular behavior that favors aggregation over degradation is contrary to what has been observed for certain thermolabile proteins, which presumably misfold upon temperature increase but are nevertheless degraded (Betting and Seufert 1996; Dohmen, Wu, and Varshavsky 1994; Downey et al. 2006). Our work instead suggests that endogenous proteins in yeast have evolved to aggregate rather than become targeted by the UPS during folding stress. This conclusion is in line with experiments demonstrating that the yeast proteome forms aggregates instead of being targeted to the UPS in response to acute heat shock (Wallace et al. 2015). Such a strategy has the advantage of preserving proteins that may be refolded at a later time after stressors are removed, allowing a faster recovery with less energy expenditure. It may also prevent adverse effects of high UPS activity, which has been shown to confer growth defects (X. Wang et al. 2010). However, by promoting aggregation, cells create a risk for proteotoxicity and entrance into a pathogenic state, as observed in the numerous neurodegenerative diseases characterized by buildup of misfolded proteins in neurons (Labbadia and Morimoto 2015; Sweeney et al. 2017). In yeast, even inert insoluble proteins can cause growth defects (Geiler-Samerotte et al. 2011). Moving forward, it remains to be determined whether human cells, particularly neurons, make a similar tradeoff that favors aggregation over degradation and whether this tradeoff puts them at risk for disease.

Because there is no surge of proteasome substrates when cells are faced with folding stressors, upregulation of the PSR in these conditions relies entirely upon the stress-sensitive transcriptional upregulation of *RPN4* rather than stabilization of the Rpn4 protein. Cells may have a divergent ability to sense different protein quality failures, as suggested by better adaptation of Cyto-Deg vs. ER-Deg under folding stress. Despite the absence of a UPS “traffic jam,” upregulation of the PSR during these stressors still improves the cell’s ability to degrade aggregation-prone proteins (Cyto-Deg and ERm-Deg in AZC and canavanine treatment). This could be because of enhanced PSR activity coinciding with the emergence of aggregates limits their formation, or because misfolded proteins are in an equilibrium between aggregated and soluble states and boosting the PSR adjusts this equilibrium point. Exploration of these possibilities is an exciting prospect for preventing and reversing diseases characterized by protein aggregation.

## Materials and Methods

### Yeast strain construction and culturing methods

All yeast strains were created from BY4741. Yeast cultures were grown at 30 °C (unless otherwise noted) in yeast extract peptone dextrose (YPD) media or synthetic dropout (SD) media.

Strain construction was done by transforming cells with a crude PCR product bearing 40 base pair overhangs homologous to the target genomic locus, a selection cassette for either antibiotic selection in YPD or auxotrophic selection in SD, and any other desired sequences. Deletion strains were constructed in the BY4741 background via transformation with antibiotic selection cassettes (NATMX6 or KANMX6), amplified with overhangs flanking the open reading frame to be deleted. The *pCYC1:RPN4* and *pYEF3:RPN4* strains were constructed via transformation by inserting a His3 cassette and 1000bp of either *pCYC1* or *pYEF3* immediately upstream of the *RPN4* open reading frame. The Hsp104-mKate2 strain was generated via transformation by inserting the yeast-optimized mKate2 coding sequence and a hygromycin selection cassette immediately downstream of the endogenous *HSP104* open reading frame. All transformants were verified by genomic PCR.

Plasmids used in this study were cloned using the Gibson Assembly method and the NEBuilder HiFi DNA Assembly Master Mix (New England Biolabs). The PSR reporter, HSR reporter, *pRPN4:GFP* reporter, Cyto-Deg, ER-Deg, and ATA-Deg plasmids were expressed from high-copy plasmids containing a Ura3 selection cassette. For *RPN4* overexpression experiments, a single-copy plasmid with a His5 selection cassette and either *pRPN4:RPN4* or no insert was co-expressed with the fluorescent reporters mentioned above. For experiments involving the expression of truncated *HSF1*, a single-copy plasmid with a Leu2 selection cassette and either *pHSF1:HSF1*_*Δ1-147*_ or no insert was co-expressed with the fluorescent reporters mentioned above.

The degron sequence for Cyto-Deg is SIFYHIGTDLWTLSEHYYEGVLSLVASVIISGR, and the degron sequence for ERm-Deg is GVKHFVFFTMFSIMPAINFPLGR (Ruggiano, Foresti, and Carvalho 2014; Maurer et al. 2016). Both sequences were connected to sfGFP by a GSGS linker.

### Fluorescent reporter assay measurements

All experiments were performed with at least three biological replicates measured on different days. For experiments testing the effects of drug stressors, stock solutions of the drugs were prepared in advance (5 mM bortezomib in ethanol, 0.5 M canavanine in water, 0.5 M AZC in water). Yeast were inoculated into selective SD media such that after overnight growth (>12 hours) in aerated culture tubes, their OD600 was between 0.05 and 0.3. Yeast were then diluted to 0.05 in a 96-well plate and incubated for 30 minutes. The drug stressors were serial diluted to 50x concentrations, then added to cells 1:50 to reach 1x concentrations. The yeast were grown for 5 hrs while shaking at 1050 rpm. Fluorescence was measured on a BD Accuri C6 flow cytometer (BD Biosciences).

Measurements of cells treated with multiple drugs (Figure 2C) differed in that drug stressors were added at two distinct timepoints according to the schematic in Figure 2c. Measurements of the knockout strains (Figure 4G) differed in that upon overnight growth, cells with OD600 between 0.05 and 0.3 were immediately measured. Comparative measurements of cells in 30 °C or 37 °C growth (Figure 4F) differed in that they were grown overnight to saturation at 30 °C, diluted to log phase and grown for 5 hours at 30 °C, then split for growth at 30 °C or 37 °C at dilutions such that they were in log phase after overnight growth, then immediately measured.

### Reporter quantification

All quantitative analysis was performed using MATLAB v8.6 (MathWorks). For the PSR, HSR, and *pRPN4:GFP* reporters, the GFP fluorescence measurements were normalized to forward scatter for each cell. For ERm-Deg, Cyto-Deg, and ATA-Deg, GFP fluorescence measurements were normalized to RFP. In experiments comparing genetic backgrounds or growth temperatures (Figure 1E, 4F, and 4G), samples were normalized to a corresponding wildtype control, which was set to 1. In titration experiments, samples were normalized to a corresponding no-treatment control that was set to 1. For titrations in backgrounds being compared to wildtype (the nonblack or broken lines in Figure 3a, 4cd, 5a), the no-treatment control was set to its mean fold value relative to the no-treatment control in solid black.

### Imaging

Cells expressing Cyto-Deg, ERm-Deg, ATA-Deg, or Hsp104-mKate2 were inoculated into selective SD media such that after overnight growth (>12 hours) in aerated culture tubes, their OD600 was between 0.1 and 0.4. Yeast were diluted to 0.1 and incubated for 30 minutes. Drug stressors were then added and the cells were incubated for 5 hours. Cells were concentrated through pelleting and resuspension, then immobilized on glass slides pre-treated with concanavalin A.

Imaging was performed on an Eclipse 80i microscope (Nikon) with an X-Cite 120LED light source (Excelitas Technologies) and using a 100x 1.40 NA oil immersion lens, controlled via MetaMorph v7.10.2.240 software (Molecular Devices). Images were captured with an Andor DR-328G-CO1-SIL Clara CCD monochrome camera (Andor Technology).

### Immunoblotting

SDS-PAGE was performed on samples using Novex Nupage 4–12% Bis-Tris gels (Thermo Fisher Scientific). Samples were then transferred onto 0.45-um nitrocellulose membranes (Bio-Rad) using a standard wet transfer protocol. Membranes were blocked with 5% milk in Tris-buffered saline with Tween for one hour at room temperature. They were then stained with 1:2,000 Pierce mouse anti-GFP (Thermo Fisher Scientific) primary antibody overnight at 4°C, followed by IRDye 800CW donkey anti-mouse (LiCor Biosciences) for three hours at room temperature. Membranes were then scanned using a LiCor Odyssey (LiCor Biosciences).

## Acknowledgements

We thank Dr. R.R. Kopito, Dr. J.E. Ferrell Jr., B.D. Alford, V. Ambati, L. Persson, C.S. Sitron, and S.L. Ergun for their helpful comments during manuscript preparation. We additionally thank Dr. J. Frydman, Dr. A. Matouschek, Dr. Y. Merbl, Dr. Z.H. Davis, and Dr. Z.A. Jaafar for helpful discussion. We thank J. Giafaglione for construction of the *pHSF1:HSF1*_*Δ1-147*_ expression plasmid and B.D. Alford for construction of the HSR reporter plasmid. We thank B.D. Alford, J. Park, and C.S. Sitron for assistance in strain building. We thank Dr. A.F. Straight and O.K. Smith for microscope usage and guidance. This work was supported by Stanford University (O.B.) and the National Institute of General Medical Sciences of the US National Institutes of Health (grant No. T32GM007276 to J.J.W.).

## Author contributions

J.J.W. and O.B. designed the experiments and wrote the manuscript. J.J.W. performed the experiments and analysis.

## References

Alford, Brian D., and Onn Brandman. 2018. “Quantification of Hsp90 Availability Reveals Differential Coupling to the Heat Shock Response.” The Journal of Cell Biology 217 (11): 3809–16.

Betting, J., and W. Seufert. 1996. “A Yeast Ubc9 Mutant Protein with Temperature-Sensitive in Vivo Function Is Subject to Conditional Proteolysis by a Ubiquitin- and Proteasome-Dependent Pathway.” The Journal of Biological Chemistry 271 (42): 25790–96.

Dohmen, R. J., P. Wu, and A. Varshavsky. 1994. “Heat-Inducible Degron: A Method for Constructing Temperature-Sensitive Mutants.” Science 263 (5151): 1273–76.

Donnelly, Michelle L. L., Martin D. Ryan, Amit Mehrotra, David Gani, Lorraine E. Hughes, Garry Luke, and Xuejun Li. 2001. “Analysis of the Aphthovirus 2A/2B Polyprotein ‘cleavage’ Mechanism Indicates Not a Proteolytic Reaction, but a Novel Translational Effect: A Putative Ribosomal ‘skip.’” Journal of General Virology. https://doi.org/10.1099/0022-1317-82-5-1013.

Downey, Michael, Rebecca Houlsworth, Laura Maringele, Adrienne Rollie, Marc Brehme, Sarah Galicia, Sandrine Guillard, et al. 2006. “A Genome-Wide Screen Identifies the Evolutionarily Conserved KEOPS Complex as a Telomere Regulator.” Cell 124 (6): 1155–68.

Ferrell, James E., Jr. 2016. “Perfect and Near-Perfect Adaptation in Cell Signaling.” Cell Systems 2 (2): 62–67.

Geiler-Samerotte, Kerry A., Michael F. Dion, Bogdan A. Budnik, Stephanie M. Wang, Daniel L. Hartl, and D. Allan Drummond. 2011. “Misfolded Proteins Impose a Dosage-Dependent Fitness Cost and Trigger a Cytosolic Unfolded Protein Response in Yeast.” Proceedings of the National Academy of Sciences of the United States of America 108 (2): 680–85.

Gidalevitz, Tali, Anat Ben-Zvi, Kim H. Ho, Heather R. Brignull, and Richard I. Morimoto. 2006. “Progressive Disruption of Cellular Protein Folding in Models of Polyglutamine Diseases.” Science 311 (5766): 1471–74.

Gidalevitz, Tali, Thomas Krupinski, Susana Garcia, and Richard I. Morimoto. 2009. “Destabilizing Protein Polymorphisms in the Genetic Background Direct Phenotypic Expression of Mutant SOD1 Toxicity.” PLoS Genetics 5 (3): e1000399.

Hahn, Ji-Sook, Daniel W. Neef, and Dennis J. Thiele. 2006. “A Stress Regulatory Network for Co-Ordinated Activation of Proteasome Expression Mediated by Yeast Heat Shock Transcription Factor.” Molecular Microbiology 60 (1): 240–51.

Ha, Seung-Wook, Donghong Ju, and Youming Xie. 2012. “The N-Terminal Domain of Rpn4 Serves as a Portable Ubiquitin-Independent Degron and Is Recognized by Specific 19S RP Subunits.” Biochemical and Biophysical Research Communications 419 (2): 226–31.

Hershko, A., H. Heller, S. Elias, and A. Ciechanover. 1983. “Components of Ubiquitin-Protein Ligase System. Resolution, Affinity Purification, and Role in Protein Breakdown.” The Journal of Biological Chemistry 258 (13): 8206–14.

Hershko, A., E. Leshinsky, D. Ganoth, and H. Heller. 1984. “ATP-Dependent Degradation of Ubiquitin-Protein Conjugates.” Proceedings of the National Academy of Sciences. https://doi.org/10.1073/pnas.81.6.1619.

Ju, Donghong, Xiaogang Wang, Haiming Xu, and Youming Xie. 2008. “Genome-Wide Analysis Identifies MYND-Domain Protein Mub1 as an Essential Factor for Rpn4 Ubiquitylation.” Molecular and Cellular Biology 28 (4): 1404–12.

Ju, Donghong, Haiming Xu, Xiaogang Wang, and Youming Xie. 2007. “Ubiquitin-Mediated Degradation of Rpn4 Is Controlled by a Phosphorylation-Dependent Ubiquitylation Signal.” Biochimica et Biophysica Acta (BBA) - Molecular Cell Research. https://doi.org/10.1016/j.bbamcr.2007.04.012.

Klaips, Courtney L., Gopal Gunanathan Jayaraj, and F. Ulrich Hartl. 2018. “Pathways of Cellular Proteostasis in Aging and Disease.” The Journal of Cell Biology. https://doi.org/10.1083/jcb.201709072.

Kraft, David Christian, Custer C. Deocaris, Renu Wadhwa, and Suresh I. S. Rattan. 2006. “Preincubation with the Proteasome Inhibitor MG-132 Enhances Proteasome Activity via the Nrf2 Transcription Factor in Aging Human Skin Fibroblasts.” Annals of the New York Academy of Sciences 1067 (May): 420–24.

Labbadia, Johnathan, and Richard I. Morimoto. 2015a. “The Biology of Proteostasis in Aging and Disease.” Annual Review of Biochemistry. https://doi.org/10.1146/annurev-biochem-060614-033955.

Lecker, Stewart H., Alfred L. Goldberg, and William E. Mitch. 2006. “Protein Degradation by the Ubiquitin–Proteasome Pathway in Normal and Disease States.” Journal of the American Society of Nephrology. https://doi.org/10.1681/asn.2006010083.

Lundgren, J., P. Masson, C. A. Realini, and P. Young. 2003. “Use of RNA Interference and Complementation To Study the Function of the Drosophila and Human 26S Proteasome Subunit S13.” Molecular and Cellular Biology. https://doi.org/10.1128/mcb.23.15.5320-5330.2003.

Ma, Menggen, and Z. Lewis Liu. 2010. “Comparative Transcriptome Profiling Analyses during the Lag Phase Uncover YAP1, PDR1, PDR3, RPN4, and HSF1 as Key Regulatory Genes in Genomic Adaptation to the Lignocellulose Derived Inhibitor HMF for Saccharomyces Cerevisiae.” BMC Genomics 11 (November): 660.

Mannhaupt, G., R. Schnall, V. Karpov, I. Vetter, and H. Feldmann. 1999. “Rpn4p Acts as a Transcription Factor by Binding to PACE, a Nonamer Box Found Upstream of 26S Proteasomal and Other Genes in Yeast.” FEBS Letters 450 (1-2): 27–34.

Maurer, Matthew J., Eric D. Spear, Allen T. Yu, Evan J. Lee, Saba Shahzad, and Susan Michaelis. 2016. “Degradation Signals for Ubiquitin-Proteasome Dependent Cytosolic Protein Quality Control (CytoQC) in Yeast.” G3 6 (7): 1853–66.

Meiners, Silke, Dirk Heyken, Andrea Weller, Antje Ludwig, Karl Stangl, Peter-M Kloetzel, and Elke Krüger. 2003. “Inhibition of Proteasome Activity Induces Concerted Expression of Proteasome Genes and de Novo Formation of Mammalian Proteasomes.” The Journal of Biological Chemistry 278 (24): 21517–25.

Moye-Rowley, W. Scott. “Transcriptional Control of Multidrug Resistance in the Yeast Saccharomyces.” Progress in Nucleic Acid Research and Molecular Biology. https://doi.org/10.1016/s0079-6603(03)01008-0.

Radhakrishnan, Senthil K., Candy S. Lee, Patrick Young, Anne Beskow, Jefferson Y. Chan, and Raymond J. Deshaies. 2010. “Transcription Factor Nrf1 Mediates the Proteasome Recovery Pathway after Proteasome Inhibition in Mammalian Cells.” Molecular Cell 38 (1): 17–28.

Ruggiano, Annamaria, Ombretta Foresti, and Pedro Carvalho. 2014. “Quality Control: ER-Associated Degradation: Protein Quality Control and beyond.” The Journal of Cell Biology 204 (6): 869–79.

Sato, Youichi, Kozue Sakamoto, Masako Sei, Ashraf A. Ewis, and Yutaka Nakahori. 2009. “Proteasome Subunits Are Regulated and Expressed in Comparable Concentrations as a Functional Cluster.” Biochemical and Biophysical Research Communications 378 (4): 795–98.

Satyal, S. H., E. Schmidt, K. Kitagawa, N. Sondheimer, S. Lindquist, J. M. Kramer, and R. I. Morimoto. 2000. “Polyglutamine Aggregates Alter Protein Folding Homeostasis in Caenorhabditis Elegans.” Proceedings of the National Academy of Sciences of the United States of America 97 (11): 5750–55.

Schmidt, Rolf M., Julia P. Schessner, Georg Hh Borner, and Sebastian Schuck. 2019. “The Proteasome Biogenesis Regulator Rpn4 Cooperates with the Unfolded Protein Response to Promote ER Stress Resistance.” eLife 8 (March). https://doi.org/10.7554/eLife.43244.

Shirozu, Ryohei, Hideki Yashiroda, and Shigeo Murata. 2015. “Identification of Minimum Rpn4-Responsive Elements in Genes Related to Proteasome Functions.” FEBS Letters 589 (8): 933–40.

Sorger, P. K. 1990. “Yeast Heat Shock Factor Contains Separable Transient and Sustained Response Transcriptional Activators.” Cell 62 (4): 793–805.

Sweeney, Patrick, Hyunsun Park, Marc Baumann, John Dunlop, Judith Frydman, Ron Kopito, Alexander McCampbell, et al. 2017. “Protein Misfolding in Neurodegenerative Diseases: Implications and Strategies.” Translational Neurodegeneration 6 (March): 6.

Szymczak, Andrea L., and Dario A. A. Vignali. 2005. “Development of 2A Peptide-Based Strategies in the Design of Multicistronic Vectors.” Expert Opinion on Biological Therapy 5 (5): 627–38.

Temple, Mark D., Gabriel G. Perrone, and Ian W. Dawes. 2005. “Complex Cellular Responses to Reactive Oxygen Species.” Trends in Cell Biology. https://doi.org/10.1016/j.tcb.2005.04.003.

Tye, Blake W., Nicoletta Commins, Lillia V. Ryazanova, Martin Wühr, Michael Springer, David Pincus, and L. Stirling Churchman. 2019. “Proteotoxicity from Aberrant Ribosome Biogenesis Compromises Cell Fitness.” eLife 8 (March). https://doi.org/10.7554/eLife.43002.

Voges, D., P. Zwickl, and W. Baumeister. 1999. “The 26S Proteasome: A Molecular Machine Designed for Controlled Proteolysis.” Annual Review of Biochemistry 68: 1015–68.

Wallace, Edward W. J., Jamie L. Kear-Scott, Evgeny V. Pilipenko, Michael H. Schwartz, Pawel R. Laskowski, Alexandra E. Rojek, Christopher D. Katanski, et al. 2015. “Reversible, Specific, Active Aggregates of Endogenous Proteins Assemble upon Heat Stress.” Cell 162 (6): 1286–98.

Wang, Li, Xicheng Mao, Donghong Ju, and Youming Xie. 2004. “Rpn4 Is a Physiological Substrate of the Ubr2 Ubiquitin Ligase.” The Journal of Biological Chemistry 279 (53): 55218–23.

Wang, Xiaogang, Haiming Xu, Seung-Wook Ha, Donghong Ju, and Youming Xie. 2010. “Proteasomal Degradation of Rpn4 in Saccharomyces Cerevisiae Is Critical for Cell Viability under Stressed Conditions.” Genetics 184 (2): 335–42.

Weids, Alan J., and Chris M. Grant. 2014. “The Yeast Peroxiredoxin Tsa1 Protects against Protein-Aggregate-Induced Oxidative Stress.” Journal of Cell Science 127 (Pt 6): 1327–35.

Xie, Y., and A. Varshavsky. 2001. “RPN4 Is a Ligand, Substrate, and Transcriptional Regulator of the 26S Proteasome: A Negative Feedback Circuit.” Proceedings of the National Academy of Sciences of the United States of America 98 (6): 3056–61.

Xu, Haiming, Donghong Ju, Tiffany Jarois, and Youming Xie. 2008. “Diminished Feedback Regulation of Proteasome Expression and Resistance to Proteasome Inhibitors in Breast Cancer Cells.” Breast Cancer Research and Treatment 107 (2): 267–74.

Yamamoto, Noritaka, Yuka Maeda, Aya Ikeda, and Hiroshi Sakurai. 2008. “Regulation of Thermotolerance by Stress-Induced Transcription Factors in Saccharomyces Cerevisiae.” Eukaryotic Cell 7 (5): 783–90.

Zheng, Xu, Joanna Krakowiak, Nikit Patel, Ali Beyzavi, Jideofor Ezike, Ahmad S. Khalil, and David Pincus. 2016. “Dynamic Control of Hsf1 during Heat Shock by a Chaperone Switch and Phosphorylation.” eLife 5 (November). https://doi.org/10.7554/eLife.18638.

